# Eosinophils are an integral component of the pulmonary granulocyte response in *Tuberculosis* and promote host resistance in mice

**DOI:** 10.1101/2021.05.29.446277

**Authors:** Andrea C. Bohrer, Ehydel Castro, Zhidong Hu, Artur T.L. Queiroz, Claire E. Tocheny, Maike Assmann, Shunsuke Sakai, Christine Nelson, Paul J. Baker, Hui Ma, Lin Wang, Wen Zilu, Elsa du Bruyn, Catherine Riou, Keith D. Kauffman, Tuberculosis Imaging Program, Ian N. Moore, Franca Del Nonno, Linda Petrone, Delia Goletti, Adrian R. Martineau, David M. Lowe, Mark R. Cronan, Robert J. Wilkinson, Clifton E. Barry, Laura E. Via, Daniel L. Barber, Amy D. Klion, Bruno B. Andrade, Yanzheng Song, Ka-Wing Wong, Katrin D. Mayer-Barber

## Abstract

Host resistance to *Mycobacterium tuberculos*is infection requires the activities of multiple leukocyte subsets, yet the roles of the different innate effector cells during tuberculosis are incompletely understood. Here we uncover an unexpected association between eosinophils and *Mtb* infection. In humans, eosinophils are decreased in the blood but enriched in resected human tuberculosis lung lesions and autopsy granulomas. Influx of eosinophils is also evident in infected zebrafish, mice, and nonhuman primate granulomas, where they are functionally activated and degranulate. Importantly, employing complementary genetic models of eosinophil deficiency, we demonstrate that, in mice, eosinophils are required for optimal pulmonary bacterial control and host survival after *Mtb* infection. Collectively, our findings uncover an unexpected recruitment of eosinophils to the infected lung tissue and a protective role for these cells in the control of *Mtb* infection in mice.

## INTRODUCTION

*Mycobacterium tuberculosis* (*Mtb*) is an intracellular bacterium that causes tuberculosis (TB), which through 2020 accounted for the highest yearly global mortality due to a single pathogen (WHO, 2020). *Mtb* primarily infects innate immune cells and host resistance depends on anti-microbial effector functions of infected macrophages alongside a robust type I and T helper 1 immune response (Cooper et al., 1993; Ernst, 2012; Flynn et al., 1993; Flynn et al., 2015). Eosinophils, in contrast, are major innate effector cells in type II inflammatory settings (Klion and Nutman, 2004; Simon et al., 2020). Accordingly, reports of type II immunity-associated cells and responses during *Mtb* infection are rare with one report of eosinophils in lungs of *Mtb*- infected guinea pigs (Lasco et al., 2004) and one case series of pulmonary eosinophilia in three TB patients (Vijayan et al., 1992). As a result, eosinophils have been assumed to be largely uninvolved in the immune response to *Mtb* or actively repressed by the strong IFNγ-driven responses required for control of mycobacteria (Cooper et al., 1993; Flynn and Ernst, 2000; Kirman et al., 2000; O’Garra et al., 2013).

Studies of eosinophil responses *in vivo* have historically focused on allergic responses and parasitic helminth infections (Klion et al., 2020; Travers and Rothenberg, 2015). Eosinophils are associated with an established type II response during lung infections with respiratory syncytial virus and *Aspergillus* (Lilly et al., 2014; Phipps et al., 2007) and recent data highlight complex roles for eosinophils in tissue homeostasis and remodeling, barrier function and wound healing (Lee et al., 2010; Rosenberg et al., 2013; Shah et al., 2020). Indeed, eosinophils are pluripotent, armed with a variety of bioactive molecules stored preformed in granules or synthesized in lipid bodies. These include a large repertoire of cytokines, chemokines, lipid mediators and cationic granule proteins including eosinophil cationic protein, eosinophil-derived neurotoxin, eosinophil peroxidase (EPX) and major basic protein (Acharya and Ackerman, 2014; Klion et al., 2020; Shamri et al., 2011). However, our understanding of the *in vivo* function of eosinophils during bacterial infections is limited. Reports are currently restricted to extracellular bacteria where direct cell-autonomous bactericidal properties of eosinophils, including extracellular DNA traps and bactericidal granule proteins, can promote extracellular clearance of *Escherichia coli*, *Citrobacter rodentium*, *Staphylococcus aureus* and *Pseudomonas aeruginosa* bacteria (Arnold et al., 2018; Krishack et al., 2019; Linch et al., 2009; Yousefi et al., 2008). However, the physiological role of eosinophils in host resistance to intracellular bacterial infections remains unknown.

Here we establish that eosinophils are a significant cellular component of TB granulomas and *Mtb*-infected lungs in mice, nonhuman primates (NHP) and patients. Using complementary genetic approaches to create eosinophil deficiency, we demonstrate that eosinophil responses are required for optimal host resistance and survival after *Mtb* infection in mice. Together our findings reveal a previously unrecognized association between eosinophils and TB in multiple hosts and uncover a protective role for these cells in mice.

## RESULTS AND DISCUSSION

We first investigated whether eosinophils may participate in the granulocytic response to TB by quantifying circulating eosinophils in three independent clinical cohort studies (Du Bruyn et al., 2020; Lowe et al., 2013; Mayer-Barber et al., 2014). We asked whether the numbers of peripheral blood eosinophils of TB patients changed based on disease status and antitubercular treatment (ATT) in a South African cohort (Du Bruyn et al., 2020). Indeed, blood eosinophil numbers were significantly higher in individuals with latent infection (LTBI) compared to active pulmonary TB (PTB) patients (**Fig. 1 A**), with a corresponding decreased eosinophil to neutrophil (E/N ratio) in the PTB group **(Fig. S1 A-B)**. When patients in an independent Chinese cohort (Mayer-Barber et al., 2014) were stratified according to disease severity based on sputum positivity for acid-fast staining of bacilli (AFB), individuals with active AFB+ PTB and extrapulmonary TB (EPTB) disease exhibited a significantly decreased E/N ratio compared to AFB- PTB patients **(Fig. S1 C)**. These data suggest that a decrease in circulating E/N ratio may reflect an increase in sputum bacterial loads. Consistent with the hypothesis that blood eosinophil numbers reflect bacterial sputum status and disease burden, circulating blood eosinophil numbers increased significantly within two weeks of commencing ATT in both cohorts **(Fig.1 B, Fig. S1 D)**. Moreover, in a third British cohort that included patients with both PTB and EPTB (Lowe et al., 2013), TB disease survivors exhibited a significantly higher E/N ratio on blood tests taken at the time of diagnosis, compared to individuals who subsequently died **(Fig. S1 E)**. In fact, lower baseline eosinophil numbers in blood correlated with shorter time to TB related death **(Fig. S1 F)**. Taken together, our data from three independent clinical cohorts suggest that peripheral blood eosinophils are decreased according to disease severity, raising the possibility that rather than being depleted, eosinophils may be recruited to distant disease sites where they could participate in the local immune response to infection.

**Figure 1:**
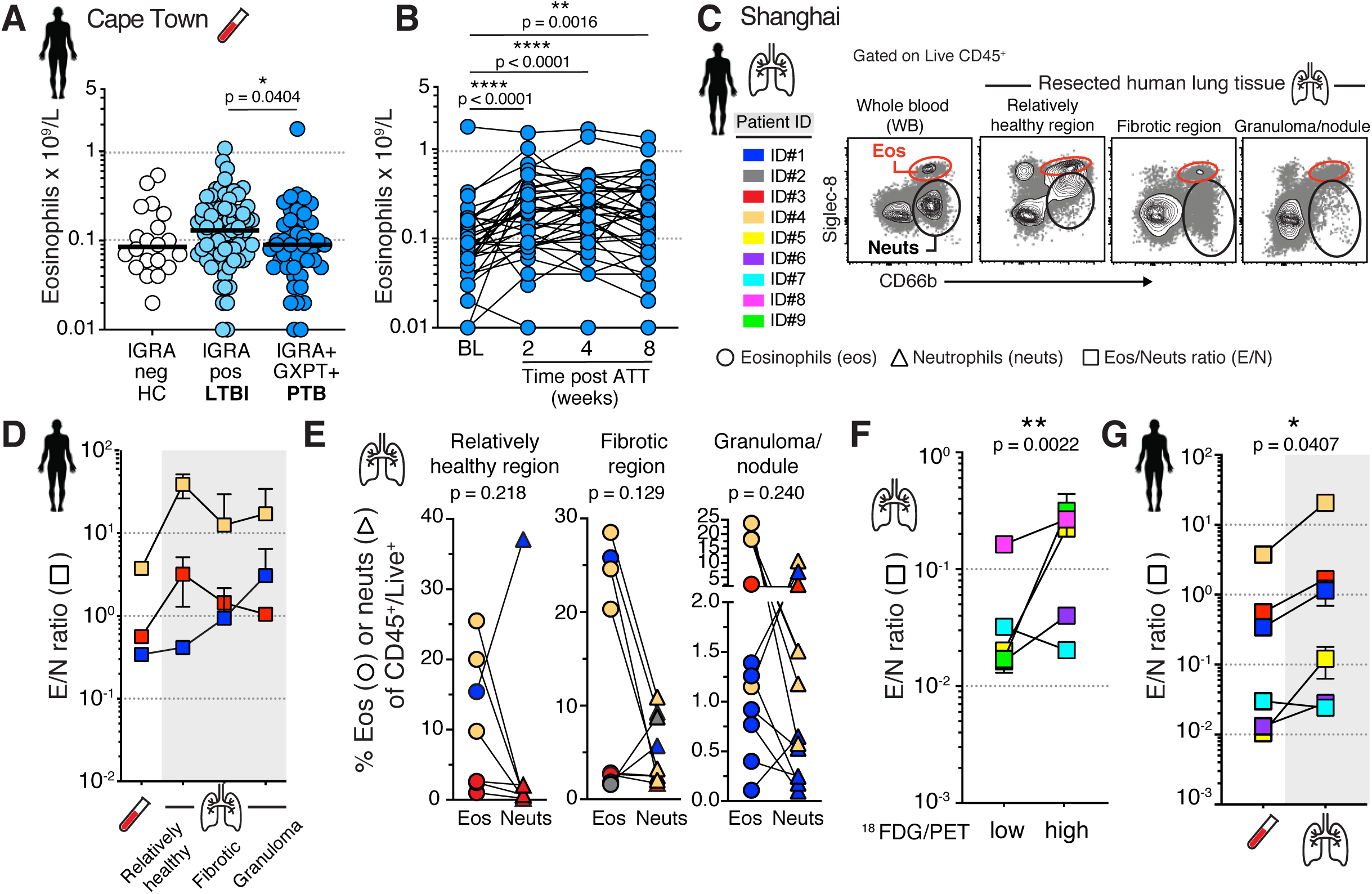
Eosinophils are decreased in circulation and enriched in human lung lesions during tuberculosis. **(A)** Cape Town cohort: Circulating eosinophil and neutrophil numbers in healthy control (HC, n=20), interferon gamma release assay (IGRA) positive latently *Mtb*-infected individuals (LTBI, n=77) and IGRA positive and GeneXpert (GXPT) positive pulmonary TB (PTB, n=48) individuals (Kruskal-Wallis with Dunn’s correction) **(B)** Cape Town Cohort: Circulating eosinophil numbers at baseline (BL) and after anti-tubercular treatment (ATT) (Wilcoxon-matched pairs test, two tailed) (n=37) **(C)** Shanghai Cohort: Representative FACS plots (from ID#1) of granulocytes in whole blood (WB) and resected human TB lesions (n=9, color coded) **(D)** Shanghai Cohort: Summary eosinophil and neutrophil ratios (E/N) in resected human lung lesions (connecting line according to color coded patient ID, tissues are depicted as mean and SEM of n=1-6 samples per tissue type and patient) **(E)** Shanghai Cohort: Eosinophil (circles) and neutrophil (triangles) proportions of CD45^+^ cells depicting individual samples per patient (connecting line, patient IDs are color coded, Wilcoxon-matched pairs test, two tailed) **(F)** Shanghai Cohort: eosinophil and neutrophil ratios (E/N) in ^18^FDG PET/CT low or high signal (SUV_max_ > 5.0) intensity lung lesions (connecting line, n=5, patient IDs color coded, Ratio paired t-test, two-tailed) **(G)** Shanghai Cohort: Summary eosinophil and neutrophil ratios (E/N) in combined resected human lung lesions (n=3=22) per patient (n=6) compared to patients’ blood eosinophils (connecting line, patient IDs color coded, Ratio paired t-test, two-tailed)

To directly address whether eosinophils could be part of the local pulmonary granulocyte response in human TB we measured the relative abundance of granulocytes in human TB lesions. To this end we generated single cell suspensions from freshly resected human lung tissue (n=9 TB patients) directly after clinically indicated surgery and identified tissue-resident granulocytes with established markers Siglec-8^+^/CD66b^+^ for eosinophils and Siglec-8^neg^/CD66b^+^ for neutrophils **(Fig. 1 C, Fig. S1 G).** Neutrophils were abundant CD45^+^ immune cells in human lung lesions divided into different macro-pathologies **(Fig. S1 H).** However, eosinophils were also frequently present in appreciable numbers relative to neutrophils in the same lung lesion sample **(Fig. 1 D, Fig. S1 H).** In fact, E/N ratios and eosinophil frequencies of CD45^+^ immune cells were sometimes similar to or even higher than neutrophils in some, but not all, samples from fibrotic areas and granulomas compared to the blood in the same individuals **(Fig. 1 D-E, Fig. S1 H-I)**. We then asked whether eosinophils would selectively infiltrate metabolically active inflamed lung lesions via 2-deoxy-2-fluorine-18-fluoro-D-glucose (^18^FDG) positron emission tomography/computed tomography (PET/CT) since the more recent patients received ^18^FDG PET/CT scans to aid in defining the surgical regions prior to resection. When we separated individual lesions based on high or low metabolic activity via ^18^FDG signal intensity, eosinophils were significantly increased relative to neutrophils in PET high compared to PET low lung lesions **(Fig. 1 F, Fig. S1 J)**. These data suggest that eosinophils are more abundant in PET FDG-avid areas compared to areas of low metabolic activity and disease. While larger studies are needed to accurately categorize and associate the abundance of specific granulocyte subsets within diverse TB lesion types, our small retrospective study showed that eosinophils can be substantially enriched in individual human TB lesions. Most importantly, the E/N ratio was significantly higher in resected human lung lesion types compared to circulation in the same patients (**Fig. 1 G**), supporting our hypothesis that peripheral blood eosinophils decrease during active TB disease and enrich at infected tissue sites. While a functional role for eosinophils in human disease remains unclear, our clinical data support the hypothesis that during TB, circulating peripheral blood eosinophils are dynamically regulated based on disease status and that eosinophils can be enriched at the *Mtb*-infected local tissue site.

To confirm the flow cytometric data, we next examined human TB granuloma sections from an independent autopsy-cohort (Blauenfeldt et al., 2018) (**Fig. 2 A**). Hematoxylin and eosin (H&E) staining indicated that eosinophils were present primarily in the rim area of the granulomas **(Fig. S2 A)**, whereas diffuse staining for the eosinophil granule protein EPX was observed in the necrotic core of granulomas, suggesting eosinophil degranulation in the center of TB granulomas (**Fig. 2 A, Fig. S2 A).** To experimentally explore the role of eosinophils, we employed a NHP model in which *Mtb* infection forms bona-fide lung granuloma structures closely resembling those found in humans (Basaraba and Hunter, 2017; Flynn et al., 2015). We first assessed lung granulomas from *Mtb*-infected macaques at 7-12 weeks post infection for eosinophil infiltration **(Fig. S2 B)**. Like the human TB granulomas, eosinophils were evident in the outer rim area in addition to more diffuse EPX staining in the necrotic core of some granulomas, suggestive of degranulated eosinophils **(Fig. 2 B, Fig. S2 C-D)**. We then asked whether eosinophil recruitment to mycobacterial granulomas was an evolutionarily conserved process. Using the zebrafish model, we assessed eosinophils in mycobacterial granulomas formed after *M. marinum* infection (Parikka et al., 2012; Swaim et al., 2006). Indeed, in agreement with our findings in NHP and human TB granulomas, eosinophils were also found in necrotic granulomas in *M. marinum* infected zebrafish (**Fig. 2 C**). Taken together, our data show that eosinophils are present in mycobacterial granulomas in human, NHP and zebrafish, suggesting that infiltration of eosinophils into granulomas may be a conserved response to mycobacteria across species.

**Figure 2:**
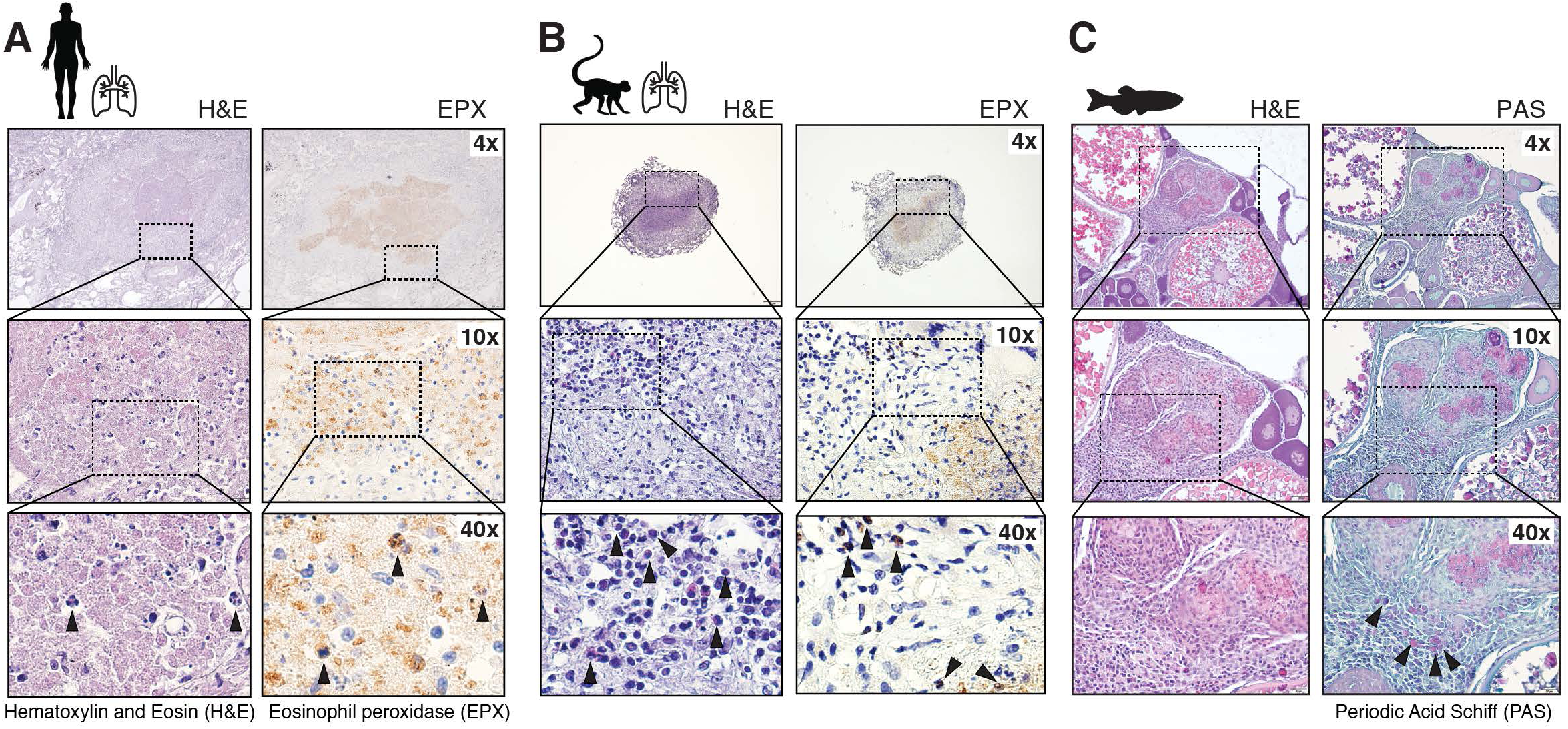
The presence of eosinophils in mycobacterial granulomas is evolutionarily conserved. (**A**) Rome cohort: Hematoxylin and Eosin (H&E) and eosinophil peroxidase (EPX) immunostaining of paraffin embedded human *Mtb* lung lesions; arrows indicate eosinophils. (**B**) H&E and EPX immunostaining of paraffin embedded rhesus macaque *Mtb* granulomas, arrows indicate eosinophils. (**C**) Organized core of a multifocal granuloma in the ovary of an *M. marinum* infected zebrafish. Black arrowheads indicate individual eosinophils stained by Periodic-acid-Schiff (PAS) in the rim of the granuloma.

To measure eosinophil activation and infiltration into granulomas we examined NHP granulomas in greater detail. In order to quantify and functionally assess eosinophils at the single cell level in rhesus macaques, we developed a flow cytometric approach based on EPX (**Fig. S2 E**). We found that intracellular staining with EPX selectively and specifically stained eosinophils in whole blood (WB), as well as in bronchoalveolar lavage fluid (BAL) (**Fig. 3 A**). Importantly, when we quantified eosinophils in NHP TB granulomas, we observed a high level of heterogeneity between individual granulomas with eosinophils spanning four orders of magnitude, comprising up to 10% of all CD45+ immune cells with an average of 1.5% in 46 granulomas from seven animals **(Fig. 3 B, Fig. S2 F).** There was no correlation between individual granuloma bacterial loads and eosinophils abundance (**Fig. 3 C**). Since, diffuse EPX staining in the core of human and NHP granulomas was suggestive of functional activation resulting in degranulation, we next quantified eosinophil degranulation by measuring CD63 surface expression (**Fig. 3 D**). While blood eosinophils showed little change in CD63 expression, BAL and granuloma eosinophils exhibited significantly increased CD63 expression with the highest CD63 expression in granulomas (**Fig. 3 D**). The proportion of CD63^+^ eosinophils varied widely between individual granulomas even within the same animal and was strongly negatively correlated with the abundance of eosinophils (**Fig. 3 E**), supporting the idea that eosinophil degranulation precedes cell death. Importantly, CD63 expression on eosinophils inversely correlated with bacterial burden in granulomas (**Fig. 3 F**), suggesting that functional activation of eosinophils in form of degranulation may impact bacterial growth in *Mtb* lesions. Collectively, our data from *Mtb* infected NHP lesions reveal a high degree of heterogeneity in the eosinophilic response and support the hypothesis that eosinophils migrate into TB granulomas where they degranulate and participate in the local innate immune response against *Mtb*.

**Figure 3:**
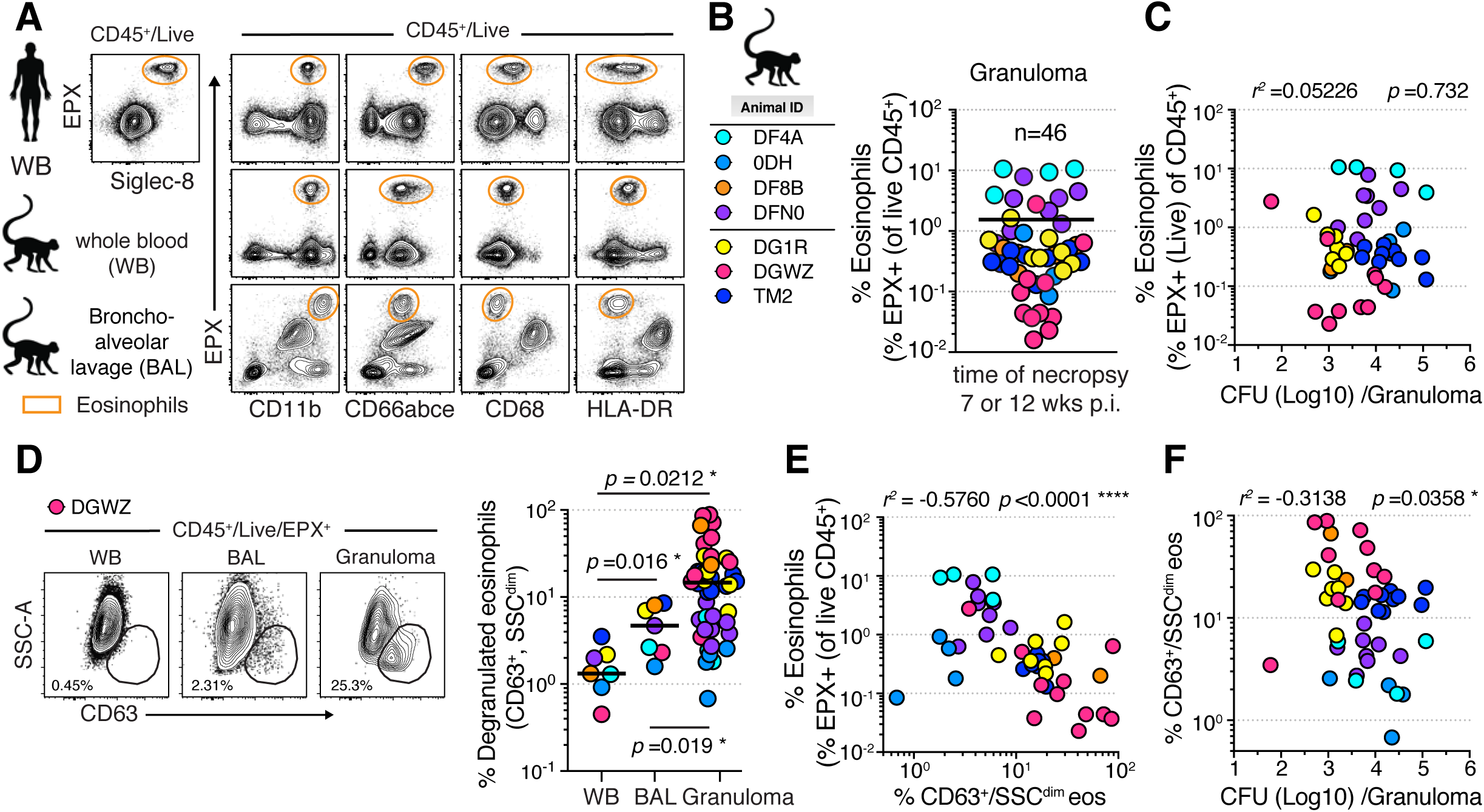
Eosinophils infiltrate and degranulate in *Mtb* granulomas of rhesus macaques. **(A)** Representative FACS plots of EPX staining of eosinophils in whole blood (WB) from healthy donors and uninfected rhesus macaques WB and bronchoalveolar lavage (BAL) **(B)** Animal ID list and corresponding color code of *Mtb* infections in present study (two independent studies, n=3-4, M+F) and percent eosinophils in pulmonary *Mtb* granulomas (n=46) **(C)** Correlation plot of eosinophil frequency and granuloma *Mtb* CFU (Spearman) **(D)** Left: Representative FACS plots of eosinophil CD63 surface expression to quantify degranulation in indicated tissues, Right: Summary data on frequency of degranulated eosinophils per individual granulomas (n=46) (Wilcoxon-Matched Pairs test for WB and BAL comparison, granuloma comparisons Ratio matched pared t-test, two tailed). **(E)** Correlation plot of frequency of CD63^+^ eosinophils and eosinophil abundance (Spearman). **(F)** Correlation plot of frequency of CD63^+^ eosinophils and granuloma *Mtb* CFU (Spearman).

As a next step in examining the function of eosinophils during *Mtb* infection we quantified eosinophils after low dose aerosol *Mtb* infection in the genetically tractable C57BL6 (B6) mouse model. We measured lung tissue resident eosinophils by flow cytometry using an intravenous (i.v.) labelling technique that distinguishes between cells located in the lung vascular capillary bed (i.v.^pos^) and cells that migrated into the lung tissue parenchyma (i.v.^neg^) (Anderson et al., 2014). When we quantified the number of lung tissue resident i.v.^neg^ eosinophils after *Mtb* infection, we observed an increase after two-three months of *Mtb* infection compared to either uninfected animals or 30 days post infection (dpi) (**Fig. 4 A, Fig. S3 A-B**). To ask whether eosinophils could be directly harboring *Mtb*, either by infection or phagocytosis, we assessed bacteria-containing eosinophils using fluorescent *Mtb*-mCherry and found that eosinophils are not infected with *Mtb* (**Fig. 4 B**). In fact, *Mtb*-infected cells have been identified and described in detail (Cohen et al., 2018; Huang et al., 2018; Lee et al., 2020; Pisu et al., 2020; Rothchild et al., 2019; Wolf et al., 2007), and we like others have consistently failed to detect significant numbers of *Mtb*-containing eosinophils (Lee et al., 2020; Rothchild et al., 2019). Thus, while eosinophils may interact with *Mtb* or *Mtb*-infected cells in lungs and TB granulomas, they themselves do not appear to represent a cellular niche for *Mtb*. Based on the increase in lung parenchymal eosinophils after infection and their role in tissue homeostasis (Lee et al., 2010; Rosenberg et al., 2013; Shah et al., 2020), we asked whether the absence of eosinophils after *Mtb* infection would lead to noticeable changes in lung pathology. To this end we infected B6 ΔdblGata mice, that lack eosinophils due to a targeted deletion of a high-affinity GATA-binding site in the GATA-1 promoter (Yu et al., 2002). However, we did not observe changes in lung gross pathology three months after *Mtb* infection in B6 ΔdblGata compared to wildtype (WT) mice (**Fig. 4 C**). We then examined lung resident immune cells and found no changes in i.v.neg *Mtb*-specific CD4^+^ and CD8^+^ T cells **(Fig. S3 C-F)**, NK1.1^+^ cells **(Fig. S3 G)**, neutrophils **(Fig. S3 H),** interstitial macrophage/dendritic cell (DC2) polarization states **(Fig. S3 I),** alveolar macrophages **(Fig. S3 J**) or XCR1+ DC1 **(Fig. S3 K)** in B6 ΔdblGata mice compared to WT mice three months after *Mtb* infection. Lastly, to explore the impact of eosinophil deficiency in an unbiased fashion, we performed whole lung transcriptional profiling on three-month *Mtb* infected B6 ΔdblGata or WT mice alongside uninfected controls (d0). Relatively few genes were significantly and differentially expressed when we directly compared transcripts from d90pi B6 ΔdblGata and d90pi WT mouse lungs. 16 differentially expressed genes (DEG) were up-regulated and nine down-regulated in *Mtb* infected ΔdblGata lungs compared to *Mtb* infected B6 control lungs **(Fig. S3 L)**. Intriguingly, most of these DEG were not associated with immunity to infection, but instead linked to neurological disorders and neuronal pathways **(Fig. S3 L,** pink annotation). To explore infection induced changes we analyzed and compared d90pi B6 ΔdblGata and d90pi WT mouse lungs to d0 WT controls. We identified 194 up- and 262 down- regulated DEG in *Mtb* infected ΔdblGata lungs, alongside 94 up- and 230 down-regulated DEG in *Mtb* infected WT B6 lungs **(Fig. S3 M)**. The top 20 DEG were again enriched in neuronal-, sensory- and olfactory-associated genes (pink annotation) in addition to genes related to lipid metabolism (green annotation) (**Fig. 4 D**). Consistent with limited changes in genes linked to host resistance, transcriptional module exploration for type I and type II interferons (Singhania et al., 2019), confirmed the restricted molecular perturbations in *Mtb* infected eosinophil deficient compared to WT lungs **(Supp. xls file)**. Importantly, gene set enrichment analyses revealed again downregulation of pathways associated with neuronal processes, neurological disorders and short chain fatty acid, endocannabinoid, and arachidonic acid metabolism in the lungs of *Mtb*- infected ΔdblGata mice **(Fig. 4E, Supp. xls file)**. Eosinophils produce a variety of bioactive arachidonic acid derived lipid mediators some of which have been implicated in host-resistance to *Mtb* infection before (Mayer-Barber and Sher, 2015). Furthermore, endocannabinoids and arachidonic acid derivatives interact with sensory neurons in the airways (Bozkurt, 2019; Chesne et al., 2019). One possible interpretation of the neuronal-associated transcriptional changes could be that they reflect alterations in airway sensory neurons or pulmonary neuroendocrine cells (PNEC) of the lung epithelium. Thus, while our transcriptional profiling of *Mtb* infected lungs revealed neuronal-associated pathways as being primarily affected by the lack of eosinophils, the biological significance of these transcriptional changes remains unknown. Moreover, the pulmonary-neuronal axis in host resistance against *Mtb* infection is largely unexplored yet there is growing appreciation for the importance of the cross talk between the nervous and immune system (Veiga-Fernandes and Mucida, 2016). For instance, a recent study found that *Mtb*-derived sulfolipids can directly bind and activate nociceptive neurons in the lung (Ruhl et al., 2020). Eosinophils have been further shown to alter parasympathetic nerve function and airway sensory nerve density thereby affecting reflex bronchoconstriction in asthma (Costello et al., 1997; Drake et al., 2018a; Drake et al., 2018b; Fryer and Wills-Karp, 1991; Kingham et al., 2002). Thus, our findings raise the possibility that eosinophils may be involved in pulmonary tissue-immune-nerve cross talk during chronic infections.

**Figure 4:**
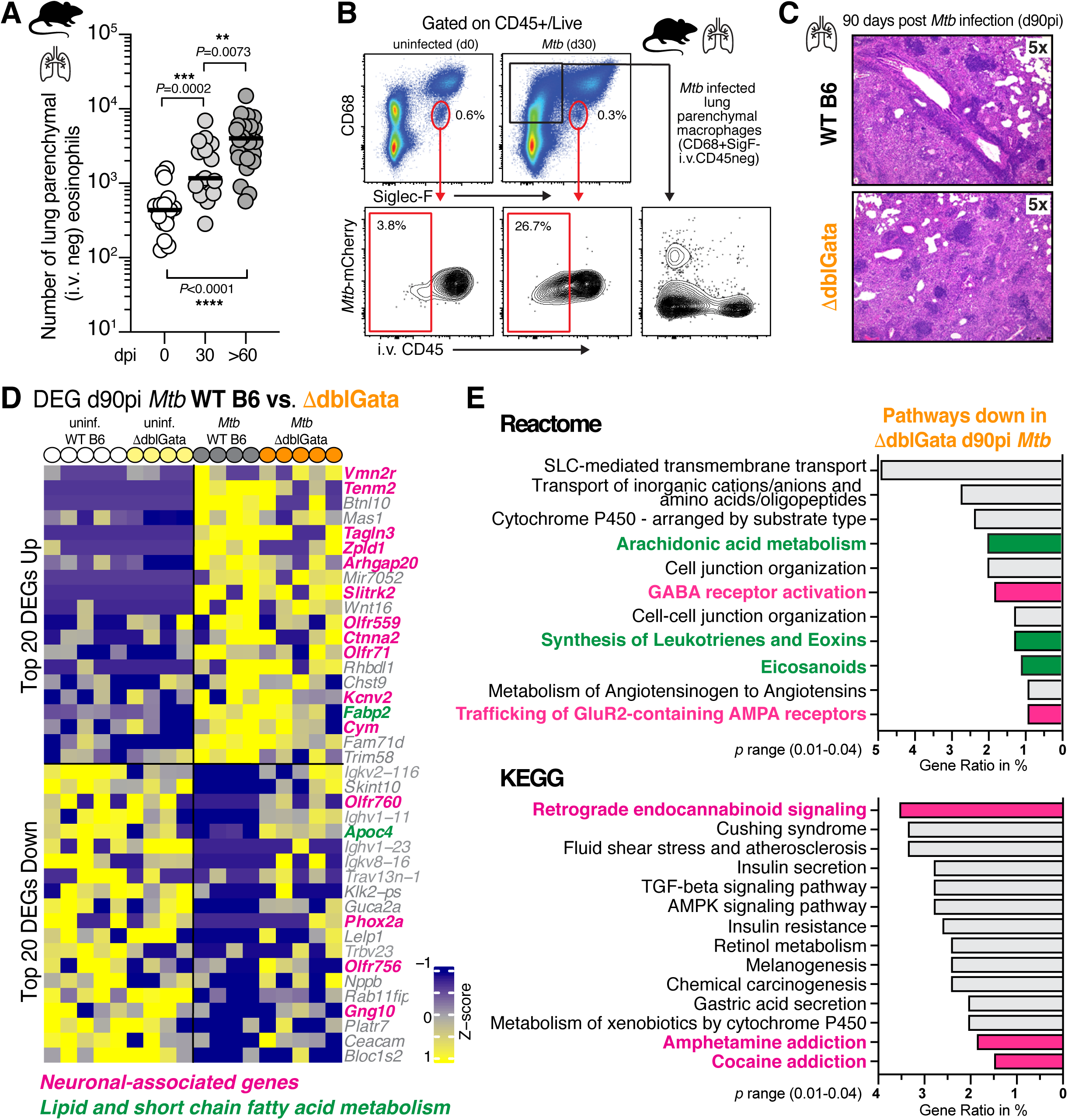
Pulmonary transcriptional profiling of *Mtb*-infected eosinophil-deficient mice reveals perturbations in lung neuronal associated pathways. **(A)** Cell number of lung parenchymal (CD45 i.v.^neg^) eosinophils over time after standard low dose (100-300 colony forming units (CFU) aerosol *Mtb* H37Rv infection in B6 mice (M+F, n=12-25 per time-point, 3-4 independent experiments, Mann-Whitney) (**B**) Representative FACS plots of intravascular (i.v.) staining and *Mtb*-mCherry quantification of eosinophils in the lungs of B6 WT mice after *Mtb* aerosol infection at d30 (3-4 independent experiments) **(C**) Representative sections of H&E staining from paraffin-embedded lungs of *Mtb*-infected (d90) WT or B6 ΔdblGata mice (M+F, n=5-6, two independent experiments) (**D**) Heatmap of top 20 up- and top 20 down- regulated DEGs of d90 WT compared to d90 B6 ΔdblGata normalized to uninfected WT mice (F, n=4-5, one experiment). Gene annotation as follows: Neuronal associated genes (pink), lipid and short chain fatty acid metabolism (green) (**E**) Gene set enrichment analysis based on Reactome (top) and KEGG (Bottom) of key pathways that are selectively downregulated in d90 B6 ΔdblGata mice. Neuronal associated pathways are highlighted in pink, lipid and short chain fatty acid metabolism pathways are highlighted in green.

Finally, to test whether eosinophils could influence the outcome of infection, we monitored pulmonary bacterial loads and survival in B6 ΔdblGata mice after *Mtb* infection. While lung bacterial loads were similar compared to WT mice for the first month, two months after *Mtb* infection we consistently measured a significant 0.5-1 log increase in the lungs of B6 ΔdblGata mice (**Fig. 5 A**). Moreover, ΔdblGata mice on either B6 or Balb/C genetic backgrounds succumbed significantly earlier to *Mtb* infection (**Fig. 5 B**). To confirm these results in a second, independent genetic model of eosinophil deficiency, we infected transgenic B6 PHIL mice that express cytocidal diphtheria toxin A under the eosinophil-specific EPX promoter (Lee et al., 2004). Like B6 ΔdblGata mice, B6 PHIL mice succumbed significantly earlier than WT mice to *Mtb* infection (**Fig. 5 C**). When infected with a high dose of *Mtb*, both B6 ΔdblGata (**Fig. 5 D**) and PHIL (**Fig. S3 N**) mice displayed significantly increased lung bacterial loads at two-three months and succumbed to *Mtb* infection-induced disease in an infectious dose-dependent manner (**Fig. 5 E-F**). Thus, the data from eosinophil-deficient mouse lines show that eosinophils play a functional role in optimal host resistance during *Mtb* infection.

**Figure 5:**
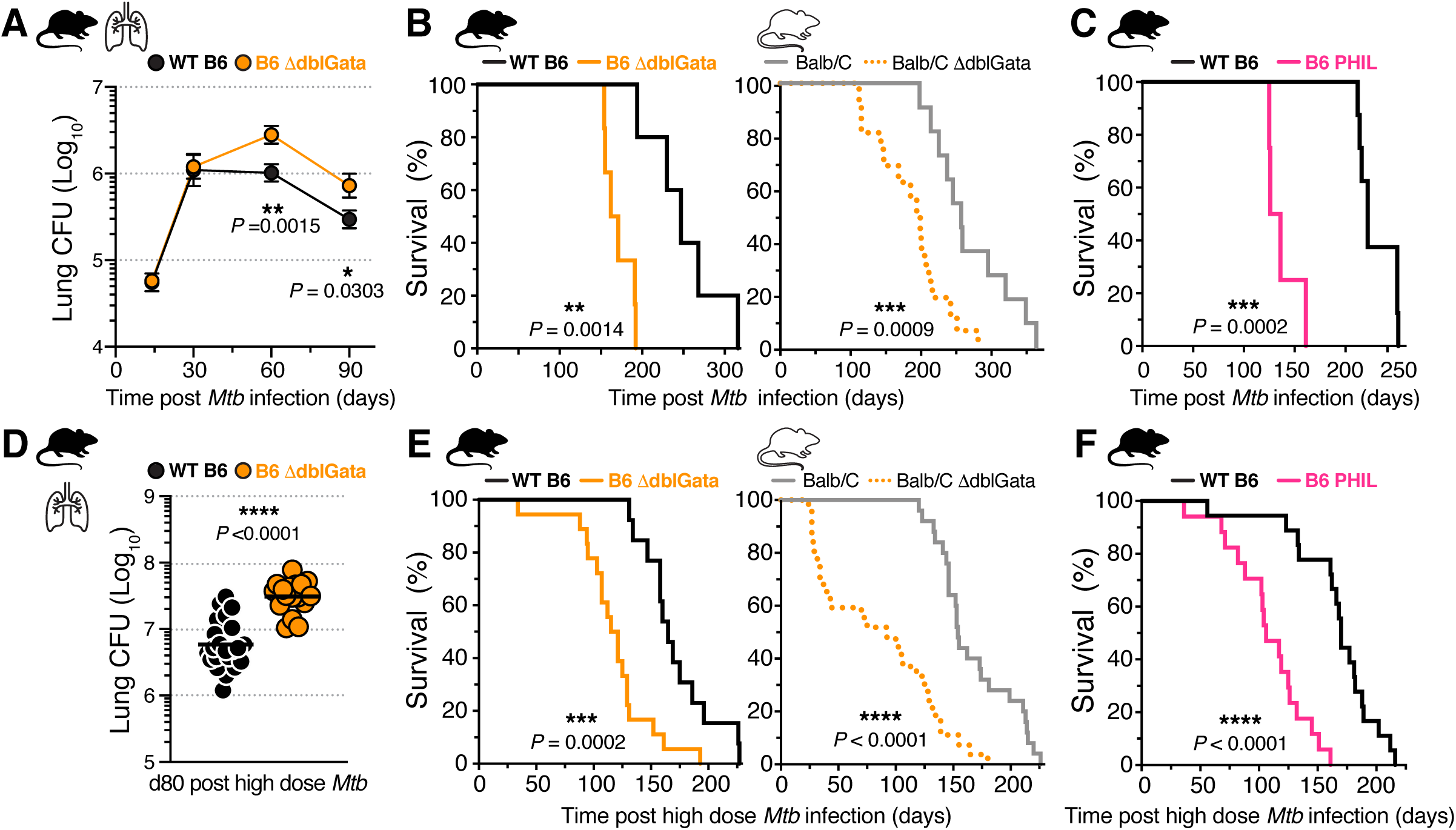
Eosinophil deficiency in mice results in increased susceptibility to *Mtb* infection. (**A**) Colony forming units (CFU) over time in lungs of standard low dose *Mtb* H37Rv (100-300 CFU) infected WT B6 or B6 ΔdblGata mice (M+F, n=11-21, 2-3 experiments per timepoint; Mann-Whitney) (**B**) Survival of WT Balb/C or B6 or ΔdblGata with LD *Mtb* (100-300 CFU) infection (M+F, LD n=5-12, 2-3 experiments each; Mantel-Cox) (**C**) Survival of WT B6 or PHIL mice after infection with 100-300 CFU *Mtb* (M+F, n=4-8, 2 experiments each; Mantel-Cox) (**D**) Lung CFU 80 days after high dose (1000-1500 CFU) *Mtb* infection of WT B6 or B6 ΔdblGata mice (M+F, n=11- 21, 3 experiments; Mann-Whitney) (**E**) Survival of WT Balb/C or B6 or ΔdblGata with 1000-1500 CFU *Mtb* infection (n= 13-27, 3-4 experiments each; Mantel-Cox).(**F**) Survival of WT B6 or PHIL mice after infection with 1000-1500 CFU *Mtb* (M+F, n=17-18, 3 experiments; Mantel-Cox).

To conclude, we found that eosinophils are recruited to *Mtb* lesions in humans, macaques, zebrafish and mice. In human lung lesions eosinophils were enriched compared to blood and found in TB granulomas. In NHP granulomas eosinophils were present and functionally activated to degranulate. We observed diffuse granule protein staining in the necrotic cores of both human and NHP granulomas, indicative of eosinophil degranulation. In settings associated with tissue necrosis *Mtb* can survive and propagate extracellularly (Basaraba and Hunter, 2017; Ernst, 2012) perhaps affording eosinophils the opportunity to access and kill extracellular *Mtb* via their characteristic degranulation and DNA trap formation. Moreover, purified human EPX has been reported to induce *Mtb* lysis *in vitro* (Borelli et al., 2003). Alternatively, granulocyte degranulation could facilitate necrotic core formation in granulomas. Thus, the described associations of eosinophils with TB granulomas in human and macaques do not allow for conclusions regarding protective or detrimental roles. In fact, it is likely that the biological function of eosinophils in host resistance to *Mtb* may be influenced by the stage and dose of infection, disease status and granulomatous features, all factors that vary considerably between the broad host species examined in this study. Nevertheless, we establish here that eosinophils represent an unanticipated integral part of the lung granulocytic response to *Mtb* across species.

Importantly, our translational approach in mice uncovered an unexpected, yet functional role of eosinophils during murine *Mtb* infection. Using multiple eosinophil-deficient mouse lines, we show that eosinophils are required for optimal host survival and bacterial control. While the underlying protective mechanisms in mice are currently unclear, they could involve anti- bactericidal or immunoregulatory effector functions that directly or indirectly affect *Mtb* growth. However, we show here that neither phagocytosis nor direct infection with *Mtb* seem likely to be occurring at significant levels in eosinophils. Additional non-direct protective mechanisms are therefore likely and could involve immunomodulatory cell-cell interactions between eosinophils and *Mtb*-infected macrophages or other pulmonary immune cells, maintenance of lung barrier function as well as disease tolerance. Such cell-cell interactions could involve production of lipid mediators, as identified in our gene set enrichment analysis, and type II cytokines. In fact, a recent study in zebrafish showed that non-canonical type II immune IL-4 and IL-13 signals played critical roles in mycobacterial granuloma formation and macrophage epithelialization (Cronan et al., 2021). Additionally, our transcriptional profiling revealed changes in neuronal associated pathways raising the intriguing possibility that eosinophils may also interact with non-immune lung cells such as PNEC and airway sensory neurons. We therefore propose that eosinophil-mediated protection against *Mtb* likely includes multiple non-mutually exclusive mechanisms, that remain to be further experimentally explored in mice and validated in interventional studies in NHPs.

Co-infection with parasitic helminths is common in areas with high TB incidence, and clinical studies have produced contradictory results on the effect of parasitic worm infections on TB outcomes (Babu and Nutman, 2016; Rafi et al., 2012). In some instances of improved clinical outcomes with TB-helminth co-infections (Abate et al., 2015; O’Shea et al., 2018; van Soelen et al., 2012) eosinophils may have contributed to improved outcomes. That said, increased susceptibility to bacterial infections has not been reported in clinical trials of eosinophil-depleting antibodies (Pavord et al., 2012; Rothenberg et al., 2008; Straumann et al., 2010) though longer-term data with examination of patients exposed to *Mtb* are needed and it is possible that a protective effect of eosinophils is masked in highly heterogenous patient populations.

Taken together, our study represents a multi-species investigation of the eosinophil response to *Mtb*. Our data reveal that eosinophilic granulocytes, typically associated with type II inflammatory responses, represent an integral part of the granulocyte response to tuberculosis, a disease associated with type I immunity. Thus, our findings open up multiple new lines of investigation into the functional relevance of eosinophils during chronic bacterial lung infection and potential new targets for host-directed therapies for tuberculosis.

## ACKNOWLEDGEMENTS

The authors thank the clinical study participants and medical staff at the Khayelitsha Site B CHC in Cape Town, the SPHCC affiliated with Fudan University in Shanghai and the HCH in Zhengzhou. We thank Drs. Y. Belkaid, A. O’Garra, S. P. Babu, M. Lionakis and A. Sher for data discussions and feedback on the manuscript. We are grateful to Ray Y. Chen for assistance with clinical protocol, all TBIP members, and the staff of the NIAID ABSL2 and ABSL3 facilities. This work was supported in part by the intramural research program of NIAID (K.D.M-B, D.L.B, C.E.B.3rd, L.E.V.) and the National Natural Science Foundation of China grant (#81770010 to K-W.W.) R.J.W is supported by NIH (U01AI115940), Wellcome (104803, 203135) and FC0010218 (CRUK, UKRI and Wellcome).

## METHODS

### Clinical study population in Shanghai, CN

Human lung resection material was obtained at the Shanghai Public Health Clinical Center (SPHCC) between 2015 and 2019 following approval by the SPHCC Ethics Committee (# 2015- S046-02 and 2019-S009-02). All patients provided written informed consent. Clinically indicated lung or pleural tissue resection surgery was performed at SPHCC as an adjunctive therapy for TB (L.W, Y.S.). Patients with HIV, other ongoing infections or immunodeficiency conditions were excluded. Using sterile instruments, resected lung tissue samples from participants ID#1-4 were divided based on macroscopically apparent diverse tissue pathologies, and a minimum of triplicate samples for each pathologic subtype were used for cell isolation and flow cytometric analysis. Participants ID#5-9 consented to ^18^FDG PET/CT signal scanning 1-6 days prior to clinically indicated surgery to guide resection and minimize risk of postoperative complications, such as increasing bronchial stump and lung resection margin embedding. After resection lung samples were further dissected based on standardized uptake values (SUV) into maximum SUV (PET high, SUV_max_ > 5.0) and low (PET low, similar to background SUV) signal pieces according to mapped 18FDG PET/CT scanning results using sterile scissors. Triplicate samples were generated for cell isolation and flow cytometric analysis for PET high and low regions from the same individuals. FACS studies were performed onsite by K.D.M-B, Z.H. and K-W.W. and samples were acquired on an LSRFortessa (BD Biosciences) and analyzed with FlowJo 10 (BD Biosciences, FlowJo, LLC).

ID#1: 28 yr. old non-smoker male, sputum smear and culture negative, received repeated ATT for 8 years, drug resistance was not detected. Treatment before surgery: isoniazid, rifampicin, pyrazinamide, and ethambutol. Surgical indication: presence of cavities and irreversible TB progression with lung destruction. Postoperative pathologic diagnosis: TB with large cavity with aspergilloma, extensive fibrotic lesions with a mixture of local cavities, granulomas, consolidations, and caseous necrosis. Acid-fast staining was 4+ positive and culture was positive for *Mtb*.

ID#2: 22 yr. old non-smoker male, sputum smear and culture negative, drug resistance was not detected. Treatment before surgery: four-month treatment with isoniazid, rifampicin, pyrazinamide, ethambutol, and levofloxacin. Surgical indication: enlargement of pleural empyema. Postoperative pathologic diagnosis: presence of inflammatory cells, fibrosis, coagulative necrosis, caseous necrosis. Acid-fast smear staining was 2+ positive and culture was positive for *Mtb*.

ID#3: 25 yr. old non-smoker female; sputum smear and culture negative at time of surgery; sputum and culture confirmed diagnosis of pulmonary tuberculosis in both lungs in 2010, repeated ATT over 6 years; drug resistance to streptomycin, isoniazid, levofloxacin and para-aminosalicylic acid were detected. Despite isoniazid, rifampicin, pyrazinamide and ethambutol regimen treatment for 1 year re-examination of chest CT showed bilateral lesions were found to be progressing and sputum culture was positive. Treatment before surgery: cycloserine, amikacin, linezolid, clofazimine, moxifloxacin, isoniazid, rifampicin, pyrazinamide, and ethambutol. Surgical indication: the presence of multi-drug resistance and cavities in the lower right lobe of the lung with complication of pneumothorax. Postoperative pathologic diagnosis: tissue infiltration of inflammatory cells, epithelioid granuloma formation, caseous necrosis and acid-fast staining (4+), which was not culture positive.

ID#4: 32 yr. old non-smoker female, 5 years of ATT with extended drug resistance to streptomycin, isoniazid, rifampicin, and ofloxacin; sputum was 4+ smear-positive and culture positive. Surgical indication: presence of extended drug resistance and consolidations and atelectasis in the lower right lobe of the lung. Postoperative pathologic diagnosis: inflammatory cell infiltration, epithelioid granuloma formation, caseous necrosis and acid-fast staining (2+), which was culture positive for *Mtb*.

ID#5: 20 yr. old non-smoker female, drug resistance was not detected, positive sputum smear, anti-TB antibody and T-spot. Sputum smear positive in July 2018 and treatment before surgery: 7-month ATT with isoniazid, rifampicin, pyrazinamide and ethambutol. Surgical indication: presence of tuberculous pleurisy related tumor with increasing size over preceding 3 months Postoperative pathologic diagnosis: inflammatory cell infiltration, epithelioid granuloma formation and coagulation necrosis. Acid-fast smear staining (4+)

ID#6: 38 yr. old male smoker (15 yrs. history), sputum smear and culture negative at time of surgery, no drug resistance, nine years prior to surgery received a one-year ATT with isoniazid, rifampicin, pyrazinamide and ethambutol after acid-fast smear positive with CT showing improved TB lesions after treatment. Symptoms relapsed three years prior to surgery, and he received ATT with isoniazid, rifampicin, pyrazinamide, ethambutol, levofloxacin, azithromycin. Surgical indication: presence of repeated TB-related symptoms including tuberculous pleuritis and cavities. Postoperative pathologic diagnosis: epithelioid granuloma and coagulative necrosis. Tissue sample acid-fast smear staining was negative.

ID#7: 39 yr. old male smoker (20 yr. history) with 5 yr. history of TB; no drug resistance, sputum smear negative at time of surgery. Previously treated with isoniazid, rifampicin, pyrazinamide and ethambutol regimen with positive sputum smear in local hospital in August 2014 for 1 year, tuberculosis recurred in June 2016, and re-anti-tuberculosis treatment for 1 year; in April 2018, the patient developed hemoptysis and chest CT suggested left upper lobe tuberculosis with cavities. He received a third anti-tuberculosis treatment. Treatment before surgery: 15-month ATT with isoniazid, rifampicin, pyrazinamide and ethambutol TB-Ab-IgG was positive. Surgical indication: clinical diagnosis of tuberculous and persistent TB cavity in spite of ATT. Postoperative pathologic diagnosis: tissue infiltration of inflammatory cells, granuloma formation, coagulative necrosis, calcification, focal pulmonary hemorrhage and local fungal infection. Acid-fast smear staining was negative and Periodic-acid-Schiff (PAS) staining was positive.

ID#8: 57 yr. old non-smoker male, sputum smear and anti-TB antibody were positive, rifampicin resistance was detected by using GeneXpert (Cepheid, Sunnyvale, CA). Treatment before surgery: 1-month ATT with rifampicin, pyrazinamide, ethambutol, isoniazid and levofloxacin (stopped due to cutaneous pruritus) and 1-month treatment with rifapentine. Surgical indication: presence of lung cavity. Postoperative pathologic diagnosis: inflammatory cell infiltration, epithelioid granuloma formation and coagulation necrosis. Fluid from postoperative lesion was acid-fast smear 3+, and GXPT-positive for *Mtb* and rifampicin sensitive.

ID#9: 23 yr. old non-smoker female, diagnosed with pulmonary tuberculosis in 2016 (positive sputum smear), isoniazid, rifampicin, pyrazinamide and ethambutol anti-tuberculosis treatment was discontinued after 6 months, fatigue, night sweats, and large pleural effusion occurred in 2018, and the sputum smear was positive again and no drug resistance prior ATT. Treatment before surgery: 10-month ATT with isoniazid, rifampicin, ethambutol, levofloxacin, bicyclol and amikacin. Surgical indication: clinical diagnosis of tuberculous hydrothorax and destructive pneumophthisis. Postoperative pathologic diagnosis: tissue infiltration of inflammatory cells, epithelioid granuloma formation, fibrosis and coagulative necrosis. GXPT test on intraoperative endocrine resected tissue revealed rifampicin sensitive *Mtb* complex.

### Clinical Study populations in Cape Town, ZA

For eosinophil studies from Cape Town, ZA, participants were recruited from the Site B Community Health Centre (CHC), in Khayelitsha between March 2017 and December 2018. Some study participants are described elsewhere (Du Bruyn et al., 2020) and in the table below. Briefly, all participants were HIV-uninfected adults (age ≥ 18 yr.) and who provided written informed consent. The study was approved by the University of Cape Town Human Research Ethics Committee (HREC 050/2015) and was conducted under DMID protocol no.15-0047. Those in the active TB group (PTB, n = 48) all tested sputum GeneXpert *Mtb*/RIF (GeneXpert (GXPT), Cepheid, Sunnyvale, CA) and/or sputum *Mtb* liquid culture positive. All active TB cases were drug sensitive and had received no more than one dose of ATT at the time of baseline blood sampling. The latent TB group (LTBI, n =77) were all asymptomatic with a positive IFNγ release assay (IGRA, QuantiFERON®-TB Gold In-Tube), tested sputum GXPT *Mtb*/RIF negative and had no clinical or radiographic evidence of active TB. Similarly, the healthy control group (HC, n=20) were all asymptomatic, without history of previous TB, tested sputum GXPT *Mtb*/RIF negative, had no clinical or radiographic evidence of active TB and tested IGRA negative. Sputum GXPT *Mtb*/RIF and differential full blood count were performed by the South African National Health Laboratory Services.

| **Cape Town** | **Healthy Control (HC);**<br>**IGRA -** | **Latent TB Infection**<br>**(LTBI); IGRA+** | **Active Pulmonary TB**<br>**(PTB); Sputum GXPT +** |
| --- | --- | --- | --- |
| **N (HIV negative)** | 20 | 77 | 48 |
| **Average age (years)** | 38 | 33 | 37 |
| **Gender (F/M)** | 12/8 | 39/38 | 13/35 |

### Clinical Study population in Zhengzhou, CN

HIV uninfected individuals with symptoms indicative of active TB who were administered less than two weeks of ATT were enrolled into a natural history study to assess response to chemotherapy (NCT01071603) conducted at the Henan Chest Hospital (HCH) in Zhengzhou from 2010 to 2012. For detailed descriptions of the clinical cohort see Mayer-Barber et al. 2014 and others (Trauner et al., 2017; Vinhaes et al., 2019).

### Clinical Study population in London, UK

The clinical study population of the TB patients in London, who were retrospectively identified by database and case-note review at the King’s College Hospital and Newham University Hospital Trust, has been described in detail in Lowe et al. (Lowe et al., 2013).

### Clinical autopsy cohort from Rome, Italy

The autopsy-cohort from the National Institute for Infectious Diseases, “Lazzaro Spallanzani” (INMI) in Rome, Italy included neutral buffered-formalin fixed, paraffin-embedded autopsy lung tissue samples from patients with histologically, histochemically, or PCR-proven confirmed pulmonary TB as previously described in Blauenfeldt et al. (Blauenfeldt et al., 2018).

### Clinical Study population in Bethesda, USA

For whole blood flow cytometry or in vitro experiments on eosinophils, healthy controls from the NIH Clinical Center (Bethesda, MD) were recruited on an institutional review board-approved clinical protocol to obtain normal blood samples (NCT00001846). All participants gave written informed consent.

### Rhesus Macaques

Thirteen healthy >2-year-old rhesus macaques (male and female, tuberculin skin test negative) were received from the NIAID breeding colony on Morgan Island. Animals were housed in nonhuman primate biocontainment racks and maintained in accordance with the Animal Welfare Act, the Guide for the Care and Use of Laboratory Animals and all applicable regulations, standards, and policies in a fully AAALAC International accredited Animal Biosafety Level 3 vivarium. All procedures were performed utilizing appropriate anesthetics as listed in the NIAID DIR Animal Care and Use Committee approved animal study proposal LPD25E. Euthanasia methods were consistent with the AVMA Guidelines on Euthanasia and endpoint criteria listed in the NIAID DIR Animal Care and Use Committee approved animal study proposal.

### *Mtb* infections of rhesus macaques

Rhesus macaques from two independent infections provided tissues for this study (Fig. S4 B) and animals DF4A, 0DH, DF8B, DFNO have previously been reported in another study (Sallin et al., 2018). Individual macaques were infected with 30-50 colony forming units (CFU) of either H37Rv or mCherry-expressing Erdman strain of *Mtb*. For infections, animals were anesthetized and 2ml of *Mtb* containing saline were instilled bronchoscopically into the right lower lung lobe. Delivered infection doses were confirmed by plating aliquots of instillation solution onto 7H11 agar plates. Colony counts were determined after 21 days.

### Mice

C57BL/6 (B6) mice were purchased from Taconic Farms (Hudson, NY). B6 PHIL mice (Lee et al., 2004) were obtained from Elizabeth Jacobsen (Mayo Clinic, Pheonix, AZ). B6 ΔdblGata (JAX 33551, B6.129S1(C)-Gata1tm6Sho/LvtzJ) mice were provided by Helene Rosenberg (NIAID). ΔdblGata (JAX 5653, C.129S1(B6)-Gata1tm6Sho/J) on the BALB/cJ background mice were purchased from Jackson Laboratories (Bar Harbor, ME). Both male and female mice, 8-24 wk. old at the onset of experiments, were used, and experimental groups in individual experiments were age and sex matched. All animals were bred and maintained in an AAALAC-accredited ABSL2 or ABSL3 facility at the NIH, and experiments were performed in compliance with an animal study proposal approved by the NIAID Animal Care and Use Committee (protocol LCIM17E).

### *Mtb* infections of mice

For aerosol infection of mice with H37Rv strains of *Mtb*, animals were placed in a whole-body inhalation exposure system (Glas-Col, Terre Haute, IN) and exposed to aerosolized *Mtb*. Delivery doses (100-300 CFU for standard low dose, 1000-1500 CFU for high dose) were confirmed by measuring lung bacterial loads at 2-24 hrs post-exposure in control mice. Bacterial loads were measured in tissue homogenates obtained via digestion and dissociation using GentleMacs (Miltenyi Biotec, CA) or mechanical homogenization using Precellys Evolution (Precellys, Atkinson, NH). Lung homogenates were serially diluted in PBS/Tween-20 and cultured on Middlebrook 7H11 agar plates supplemented with oleic acid-albumin-dextrose-catalase (Difco, Detroit, MI). Colony counts were determined after 21 days. H37Rv-mCherry was provided by Kevin Urdahl (Cohen et al., 2018).

### *M. marinum* infection and paraffin embedding of zebrafish

All zebrafish husbandry and experiments were approved by the Duke University Animal Care and Use Committee (protocol A122-17-05). Single cell aliquots of cerulean-expressing *M. marinum* were made according to the method of Takaki (Takaki et al., 2013). Zebrafish were anesthetized with 0.016% tricaine and infected with *M. marinum*-expressing cerulean fluorescent protein at a dose of 400 fluorescent bacteria/animal by intraperitoneal injection of the zebrafish. Zebrafish were maintained in spawning tanks with daily water changes. At 2 weeks post infection, zebrafish were euthanized by tricaine overdose and the head and the tail of the zebrafish were removed with a scalpel. A small incision was made in the belly of the zebrafish and the zebrafish was fixed for 2 days in 4% paraformaldehyde. The animals were subsequently processed and paraffin-embedded by the Duke Research Immunohistology Laboratory.

### Histopathology

Tissue samples were fixed with 4% paraformaldehyde, paraffin-embedded, sectioned and stained with either hematoxylin and eosin (H&E), the Ziehl–Nielsen histochemical method to visualize acid-fast mycobacteria, anti-EPX for the immunohistochemical identification of eosinophils or Periodic-acid-Schiff (PAS) for necrosis. Eosinophils were identified via light microscopy using an Olympus BX51 microscope (magnifications of 4x, 10x or 40x) and photomicrographs were taken using an Olympus DP73 camera. The distribution of eosinophil staining was assessed microscopically using indicated scales; all histological analyses were scored in a blinded manner by a board-certified veterinary pathologist.

### Confocal microscopy of rhesus macaque granulomas

Excised granulomas from Erdman-mCherry fluorescent *Mtb*-infected rhesus macaques were fixed overnight in 4% paraformaldehyde and subsequently embedded in Optimal cutting temperature compound (OCT, Fisher Scientific) and stored at -80°C . Cryostat cut sections (7μm) were placed onto microscope slides (Fisher Scientific) and sections were blocked with 10% FCS in PBS and incubated with anti-Laminin-AF647 and anti-human EPX AF488 for 1h incubation at 37°C in a humidified chamber. Afterwards the slides were washed in PBS for 10 min and mounted with ProLong® Gold Antifade mountant (ThermoFisher). The samples were analyzed with a Leica SP8 confocal laser fluorescence microscope and Imaris software (Bitplane).

### RNA extraction and RNA-Seq

For RNA extraction dedicated lung lobes were placed in RNAlater (Invitrogen, Carlsbad, CA) and stored at -80°C. RNAlater-stabilized lung lobes were thawed at RT for 20min, then homogenized in RLT plus buffer with β-mercaptoethanol (Qiagen, Hilden, Germany). Total RNA was then isolated from the RLT-homogenized cells using the RNeasy Plus Mini Kit (Qiagen, Hilden, Germany). RNA quantification, qualification as well as library preparation and transcriptome sequencing was performed by Novogene (Novogene Corporation Inc, Sacramento, CA). Briefly, RNA degradation and contamination was monitored on 1% agarose gels and RNA purity was checked using the NanoPhotometer® spectrophotometer (IMPLEN, CA, USA). RNA integrity and quantitation were assessed using the RNA Nano 6000 Assay Kit of the Bioanalyzer 2100 system (Agilent Technologies, CA, USA). A total amount of 1 μg RNA per sample was used as input material for the RNA sample preparations. Sequencing libraries were generated using NEBNext® UltraTM RNA Library Prep Kit for Illumina® (NEB, USA) following manufacturer’s recommendations and index codes were added to attribute sequences to each sample. Briefly, mRNA was purified from total RNA using poly-T oligo-attached magnetic beads. Fragmentation was carried out using divalent cations under elevated temperature in NEBNext First Strand Synthesis Reaction Buffer (5X). First strand cDNA was synthesized using random hexamer primer and M-MuLV Reverse Transcriptase (RNase H-). Second strand cDNA synthesis was subsequently performed using DNA Polymerase I and RNase H. Remaining overhangs were converted into blunt ends via exonuclease/polymerase activities. After adenylation of 3’ ends of DNA fragments, NEBNext Adaptor with hairpin loop structure were ligated to prepare for hybridization. In order to select cDNA fragments of preferentially 150∼200 bp in length, the library fragments were purified with AMPure XP system (Beckman Coulter, Beverly, USA). Then 3 μl USER Enzyme (NEB, USA) was used with size-selected, adaptor ligated cDNA at 37 °C for 15 min followed by 5 min at 95 °C before PCR. Then PCR was performed with Phusion High-Fidelity DNA polymerase, Universal PCR primers and Index (X) Primer. At last, PCR products were purified (AMPure XP system) and library quality was assessed on the Agilent Bioanalyzer 2100 system. The clustering of the index-coded samples was performed on a cBot Cluster Generation System using PE Cluster Kit cBot-HS (Illumina) according to the manufacturer’s instructions. After cluster generation, the library preparations were sequenced on an Illumina Novaseq 6000 platform and paired-end 150bp reads were generated. The entire gene expression data set will be available at the GEO database accession number GSE165871.

### Transcriptional analysis

For all samples, low-quality bases were removed and adapters were trimmed using *Trimmomatic V0.32* (Bolger et al., 2014). After the quality check, sequences were aligned to the *Mus musculus* genome (GRCm38.p6 version 67), with *STAR* 2.7.0 (Dobin et al., 2013). After mapping, the output was converted to count tables with the *tximport* package (Soneson et al., 2015) from *R 4.0.2*. The stratification was performed as follows: d0 WT B6 group was composed of 5 samples: d0WT1- d0WT5, d0 ΔdblGata group (d0Gata1-4), d90 WT B6 group (d90WT1-4) and d90 ΔdblGata group (d90Gata1-5). Count gene expression matrix was examined by *DESeq2* packed (Love et al., 2014) from *R 4.0.2* to identify differentially expressed genes (DEG) following the comparisons: d90 ΔdblGata vs. d90 WT B6, d0 ΔdblGata vs. d0 WT B6, d90 ΔdblGata vs. d0 WT B6, d90 WT B6 vs. d0 WT B6. Changes in gene expression levels were considered significant when statistical test values (FDR adjusted p-value) were lower than 0.05 and the fold change/difference higher than ±1.4. Candidate DEG were visualized in volcano plots and Venn diagrams with *EnhancedVolcano* and *VennDiagram* packages from *R 4.0.2*. The obtained DEG were scanned by *REACTOME* (Yu and He, 2016) and KEGG (http://www.genome.jp/kegg/) pathway database using *compareCluster* package (Yu et al., 2012) from *R 4.0.2*. The gene list of interferon-inducible (Type I) and IFN-γ-inducible (Type II) lung modules were retrieved from Singhania *et al*. (Singhania et al., 2019). The sample clustering and classification were assessed using Heatmaps, applied in the variance stabilizing transformation gene expression values.

### Cell Isolations for flow cytometry - Peripheral blood

Rhesus macaque and human blood samples were collected in EDTA tubes and whole blood was used for flow cytometry. Briefly, antibody-cocktails were directly added to 200μl of whole blood and stained at 37°C for 20min, after which samples were washed twice in 4ml 1% FCS/PBS. Fixable live/dead cell stain (Molecular Probes-Invitrogen) was used according to the manufacturer’s protocol. Pellets were then fixed and permeabilized with the Foxp3 Transcription Factor Staining Buffer Kit (Life Technologies/eBioscience) for at least 1hr followed by intracellular antigen staining for 30min at 4°C. Cells were washed and samples acquired on a FACSymphony (BD Biosciences) at NIH or a LSRFortessa (BD Biosciences) at SPHCC. FACS data were analyzed using FlowJo 10 (BD Biosciences, FlowJo, LLC).

### Cell Isolations for flow cytometry - Bronchoalveolar lavage (BAL) and granulomas from rhesus macaques

Rhesus macaque BAL samples were passed through a 100μm cell strainer, pelleted, and counted for analysis. Granulomas were individually resected from the lungs and samples used for flow cytometry analysis were pushed through a 100μm cell strainer. Aliquots from all samples were serially diluted and plated on 7H11 agar plates for CFU quantification. Alternatively, resected granulomas were fixed in 4% paraformaldehyde for later paraffin embedding. Samples were stained with surface antibody-cocktails for 20min at 4°C, followed by fixable live/dead cell stain (Molecular Probes-Invitrogen). Cells were then fixed and permeabilized (eBioscience Transcription Factor Staining Buffer Kit) for at least 1hr followed by intracellular antigen staining for 30min at 4°C. Cells were washed and samples acquired on a FACSymphony (BD Biosciences) at NIH. FACS data were analyzed using FlowJo 10 (BD Biosciences, FlowJo, LLC).

### Cell Isolations for flow cytometry - Human and murine lung tissue

Mice were intravenously (i.v.) injected 3 min. prior to euthanasia with 5μg/mouse of APC or BV711 labelled CD45 (30-F11) as previously reported (Anderson et al., 2014). Lungs from infected mice were digested using the Miltenyi lung cell isolation buffer and dissociated via GentleMACS (Miltenyi Biotec, CA) according to the manufacturer’s instructions. Digested lung was fully dispersed by passage through a 100µm pore size cell strainer and an aliquot was removed for bacterial load measurements. Cells were then washed and purified using a 37% Percoll step. Samples were stained with surface antibody-cocktails for 20min at 4°C followed by fixable live/dead cell stain (Molecular Probes-Invitrogen). Cells were then fixed and permeabilized (eBioscience Transcription Factor Staining Buffer Kit) for at least 1hr followed by intracellular antigen staining for 30min at 4°C. Cells were washed and samples acquired on a FACSymphony (BD Biosciences) at NIH. FACS data were analyzed using FlowJo 10 (BD Biosciences, FlowJo, LLC).

The surgically-resected human lung tissues samples were weighed and manually aliquoted into replicates using sterile scissors inside a sterile 50ml tube. Aliquots were transferred into Trizol for RNA isolation or subjected to single cell isolation. Briefly, sample replicates for single cell isolation were digested with 100U/ml of Collagenase IV (Sigma), 50 U/ml of Benzonase (Sigma) in 8ml of incomplete RPMI-1640 medium (without fetal bovine serum) for 45min at 37°C in the shaker. Cells were then filtered through 100μm cell strainers (Miltenyi), and the remainder of tissue pieces were gently squashed by syringe and washed with 4ml ice-cold PBS containing 50% FBS to stop digestion. Next, the cell suspension was centrifuged and subjected to Debris Removal Solution (Miltenyi) according to the manufacturer’s protocol. Samples were stained with surface antibody- cocktails for 20min at 4°C, followed by fixable live/dead cell stain (Molecular Probes-Invitrogen). Cells were then fixed and permeabilized (eBioscience Transcription Factor Staining Buffer Kit) for at least 1hr followed by intracellular antigen staining for 30min at 4°C. Cells were washed and samples acquired on a LSRFortessa (BD Biosciences) at SPHCC. FACS data were analyzed using FlowJo 10 (BD Biosciences, FlowJo, LLC).

### Flow cytometry reagents

Fluorochrome-labeled antibodies against mouse, human or rhesus macaque antigens used for flow cytometric analysis are listed below: I -A/I-E (clone M5/114.15.2), Ly6G (1A8), CD11c (HL3 and N418), CD45.2 (104), TCRb (H57-597), NK1.1 (PK136), CD11b (M1/70), CD45 (30-F11), CD68 (FA-11), Ly6C (AL-21 and HK1.4), Siglec-F (E50-2440), CD193 (J073E5), Arg1 (A1exF5), NOS2 (CXNFT), CD4 (L3T4), CD8a (53-6.7), Foxp3 (FJK-16s), IRF8 (V3GYWCH), XCR1 (ZET), CD193 (5E8.4), EPX (AHE-1), CD62L (MEL-14), CD14 (M5E2), CD11c (3.9), CD45 (D058- 1283), CD66abce (TET2), CD15 (HI98), CD68 (Y1/82A), HLA-DR (L243), CD16 (3G8), CD45 (2D1), CD193 (83101), CD66b (G10F5), CD62L (REG-56), CD206 (15-2.2), CD123 (6H6) and Siglec-8 (7C9). ESAT-6_1–17_ or TB10.4_4–11_ tetramers were produced by the NIAID tetramer core facility (Emory University).

### Statistical analyses

The statistical significance of differences between data groups were calculated using GraphPad Prism 8 as indicated in the figure legends. The Mann-Whitney test, the Wilcoxon matched pairs test (paired samples) were used for comparison of group means, Log-rank Mantel-Cox test for survival data and Spearman correlation test unless otherwise indicated in the figure or figure legends.

### Description of supplemental material

Fig. S1 shows additional clinical data and study designs Fig. S2 human and nonhuman primate granuloma structure with eosinophil distribution and EPX staining specificity Fig. S3 shows depletion efficiency in eosinophil deficient mouse strains after *Mtb* infection, additional transcriptional and immunological parameters assessed in ΔdblGata and increased bacterial loads in B6 PHIL mice.

---

**Supplemental Figure S1:**
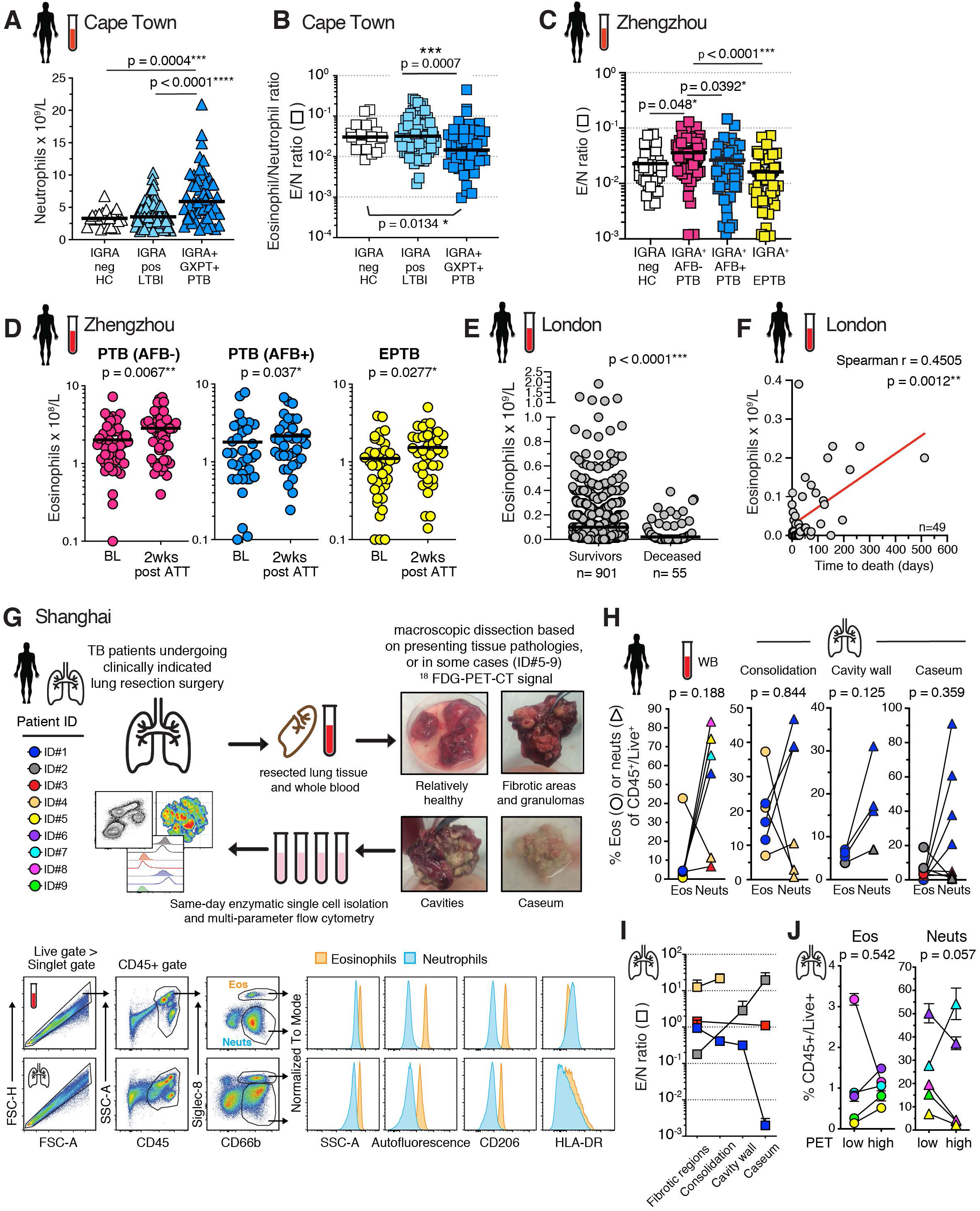
Peripheral blood eosinophils in clinical cohorts and granulocyte isolation of from human TB lung lesions. **(A)** Cape Town Cohort: Circulating neutrophil numbers and **(B)** eosinophil/neutrophil (E/N) ratio in interferon gamma release assay (IGRA) negative healthy controls (HC, n=20), IGRA positive latently *Mtb*-infected individuals (LTBI, n=77) and GeneXpert (GXPT) positive pulmonary TB (PTB, n=49) individuals (Kruskal-Wallis with Dunn’s correction) **(C)** Zhengzhou Cohort: eosinophil/neutrophil (E/N) ratio in HC (n= 30), acid fast bacilli (AFB) staining neg (AFB-, sputum negative), PTB (n=64, pink), and AFB positive (AFB+, sputum positive) PTB (n=48, blue), and extrapulmonary TB (EPTB, n=50, yellow) (Kruskal-Wallis with Dunn’s correction) **(D)** Zhengzhou Cohort: Circulating eosinophil numbers at baseline (BL) and 2 wks. after anti-tubercular treatment (ATT) (Matched Wilcoxon) from color coded clinical groups (pink, IGRA+AFB-PTB, n=47 pairs; blue, IGRA+AFB+PTB, n=34 pairs; yellow, EPTB, n=30 pairs). **(E)** London Cohort: Circulating eosinophil numbers in survivors (n=901) or deceased (n=55) TB patients (Mann-Whitney). **(F)** London Cohort: Circulating eosinophil numbers correlated with time to death (Spearman correlation, n=49). **(G)** Schematic study overview and examples of human lung TB lesions after resection surgery (n=9) and flow cytometric gating strategy of human lung TB lesions for eosinophil and neutrophil quantification in Shanghai cohort. **(H)** Shanghai Cohort: Eosinophil (circles) and neutrophil (triangles) proportions of CD45^+^ cells depicting individual samples per patient (connecting line, patient IDs are color coded, Wilcoxon-matched pairs test, two tailed) (**I**) Shanghai Cohort: Summary eosinophil and neutrophil ratios (E/N) in resected human lung lesions (connecting line according to color coded patient ID, tissues are depicted as mean and SEM of n=1-6 samples per tissue type and patient). **(J)** Shanghai (SH) Cohort: Eosinophil (circles) and neutrophil (triangles) proportions of CD45^+^ cells in ^18^FDG PET/CT low or high signal (SUV_max_ > 5.0) intensity lung lesions (connecting line, n=5, patient IDs color coded, Wilcoxon-matched pairs test, two tailed).

**Supplemental Figure S2:**
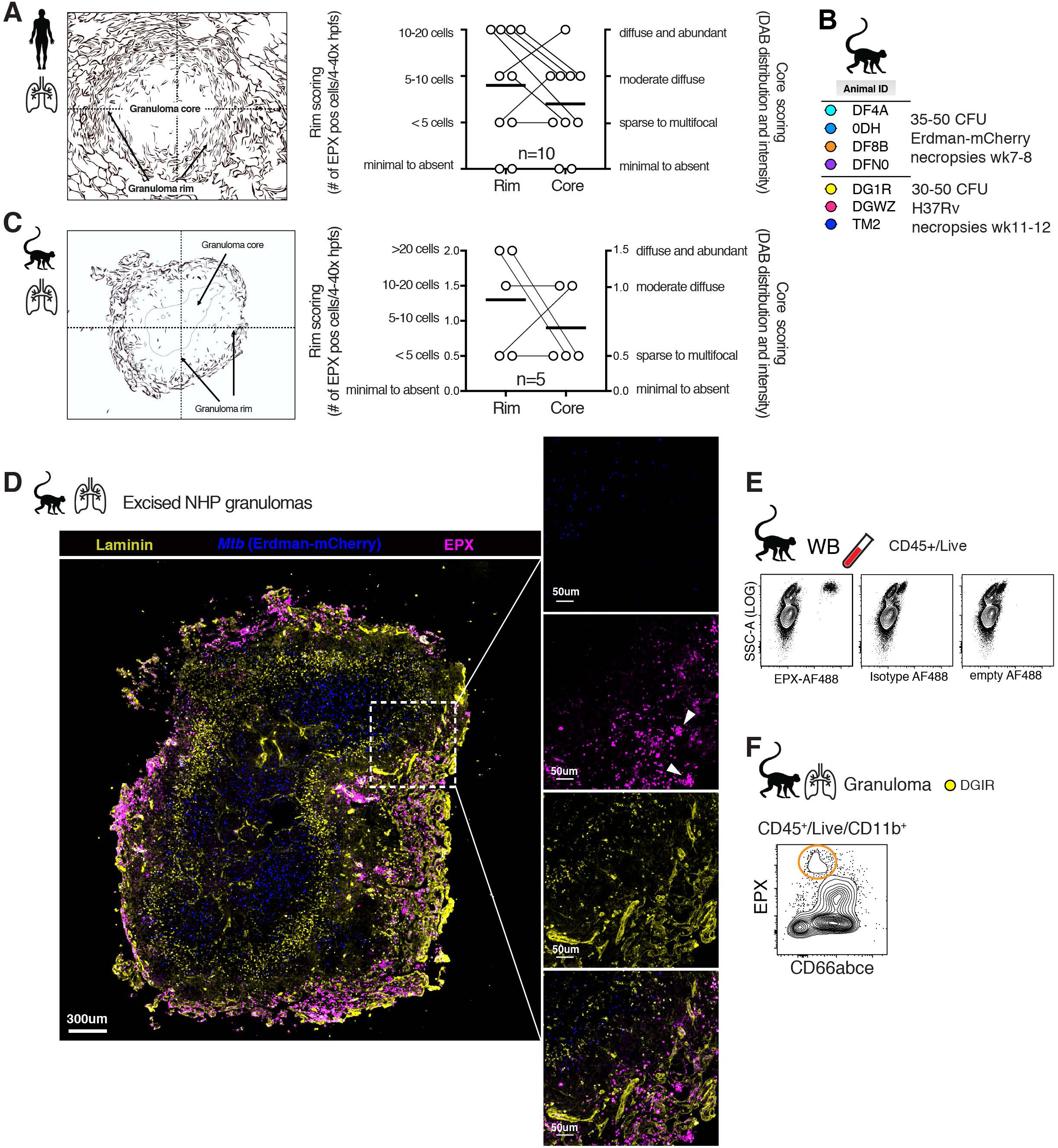
Human and NHP TB lesions assessed by H&E and EPX immunohistological and flow cytometric staining. **(A)** Rome cohort: Histological scoring of eosinophil distribution in human TB lesion on blinded specimens (n=10). **(B)** Animal IDs and infection dose and strains for NHP Mtb infections (n= 7, M+F), two independent experiments **(C)** Granuloma outline and legend for histological scoring of NHP granulomas on blinded specimens (n=5). **(D)** EPX immunofluorescence staining on thin sections from paraffin embedded granulomas after Erdman-mCherry (blue) *Mtb* infection. **(E)** Isotype and fluorescence minus one FMO (empty) control of EPX-AF488 staining in NHP whole blood (WB). **(F)** Example EPX FACS staining in granulomas from rhesus macaques.

**Supplemental Figure S3:**
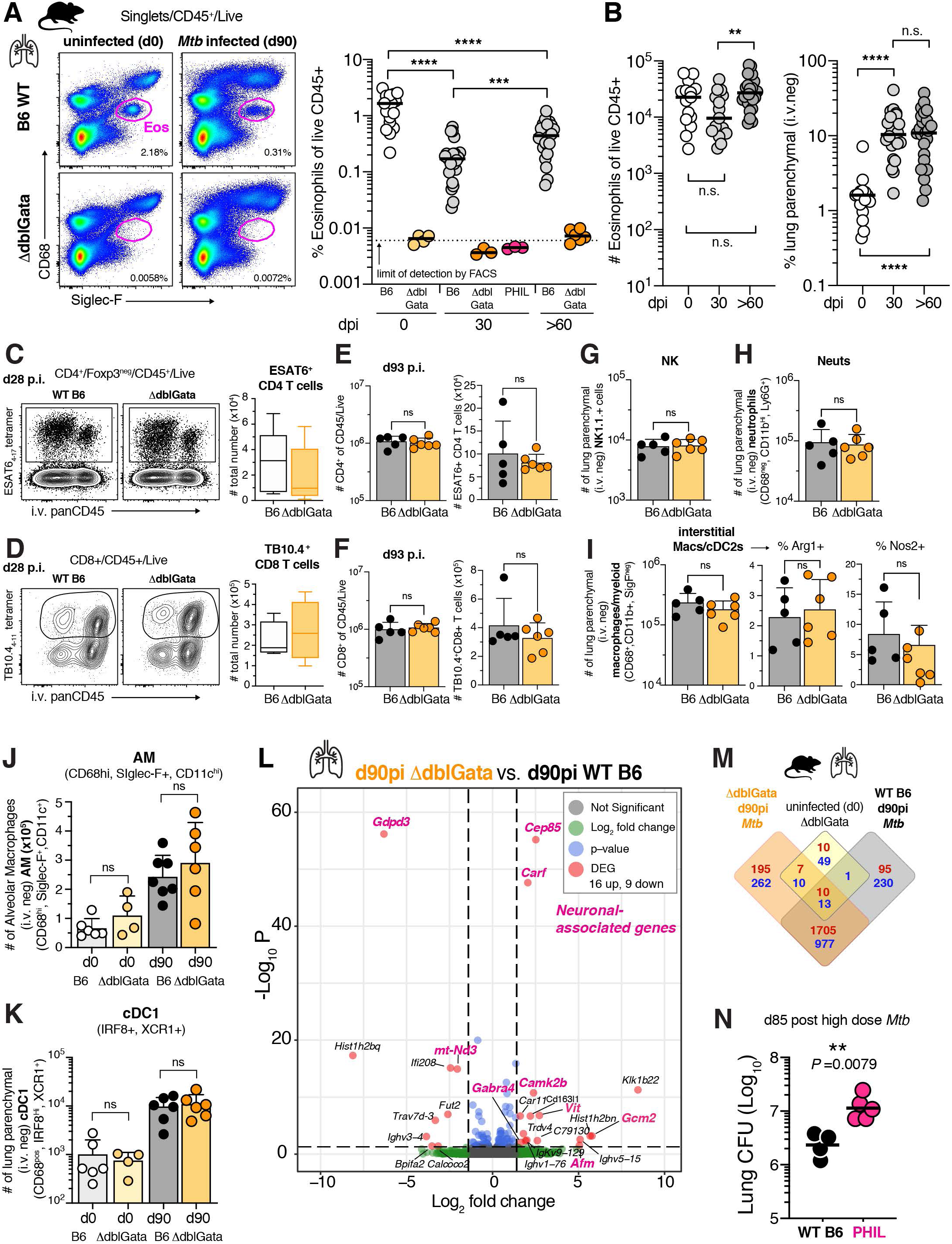
Transcriptional changes and immune profiling in eosinophil deficient mice after *Mtb* infection. (**A**) Example flow cytometry of eosinophils in B6 WT and ΔdblGata mice and lung eosinophil frequency in naïve and Mtb infected (100-300 CFU) B6 WT and eosinophil deficient mice. (M+F, n=3-28, 1-4 independent experiments per timepoint, Mann-Whitney) **(B)** Numbers of total and lung parenchymal (i.v. neg) eosinophils at indicated time points after *Mtb* infection (M+F, n=12- 25, 3-4 independent experiments per timepoint, Mann-Whitney) (**C**) Representative flow cytometry for intravascular stain and ESAT6_4-17_ CD4 tetramer with quantification of *Mtb*-specific CD4^+^ T cells and (**D**) *Mtb*-specific CD8+ T cells 4 weeks p.i. (M+F, n=8-10, Mann-Whitney, 2 experiments). Quantification of (**E**) total CD4+ cells and ESAT6_4-17_ CD4 tetramer+ CD4^+^ T cells and (**F**) total CD8+ and 10.4_4-11_ specific CD8 tetramer+ cells three months after infection (M+F, n=5-6, Mann-Whitney, 2 experiments). Quantification at three months post infection of (**G**) Nk1.1^+^ cells, (**H**) neutrophils, **(I)** interstitial macrophages (Macs)/dendritic cell 2 (DC2) and their corresponding frequency of arginase (Arg1) or iNOS (Nos2) expression (M+F, n=5-6, Mann- Whitney, 2 experiments). Quantification at baseline and three months post infection of (**J**) alveolar macrophages (AM) and (**K**) XCR1+ conventional dendritic cells 1 (cDC1) (F, n=4-6, Mann- Whitney, 1 experiment). (**L**) Volcano plot of differentially expressed genes (DEG, red) between lungs of d90 WT (n=4) or B6 ΔdblGata (n=5) *Mtb*-infected mice. Green are genes with Log2 Fold Change ±1.4, Blue are genes with a p-value and False discovery rate (FDR) lower than 0.05 and grey genes are not significant. The changes in gene expression levels were considered significant when statistical test values (FDR adjusted p-value) were lower than 0.05 and the fold change/difference higher than ±1.4. Neuronal associated genes are annotated in pink. One experiment with n=4=5 per group. (**M**) Venn diagram of DEG in lungs from three-month (d90) infected WT B6 (n=4) or B6 ΔdblGata mice (n=5) and uninfected (d0) B6 ΔdblGata (n=4) compared to lungs from uninfected (d0) WT B6 mice (n=5). Upregulated DEG are shown in red and downregulated DEG in blue. One experiment with n=4=5 per group (**N**) Lung bacterial loads 85 days after high dose *Mtb* infection in eosinophil deficient B6 PHIL mice (M, n=4-5, Mann- Whitney, 1 experiment).

